# Predicting Theory of Mind in children from the infant connectome

**DOI:** 10.1101/2024.05.22.595346

**Authors:** Clara Schüler, Philipp Berger, Charlotte Grosse Wiesmann

## Abstract

Our ability to reason about other people’s mental states, labeled Theory of Mind (ToM), is critical for successful human interaction. Despite its importance for human cognition, early predictors of individual ToM development are lacking. Here, we trained a computational model to identify whole-brain connectivity patterns predictive of joint attention, from resting-state fMRI data of 8-15-month-old infants, and tested whether the identified connectome would also predict ToM capacity later in development. First, the model significantly predicted joint attention scores in an independent infant sample. Crucially, the identified connectome did indeed predict ToM in children aged 2-5 years. The default network and its interaction with the ventral attention network formed dominant connections of the network, suggesting that the interplay of bottom-up attention and higher-order cognition paves the way for mature social cognition. These findings provide an early marker for individual differences in social cognitive development, with high potential for the early diagnosis of social cognitive disorders.

---

As inherently social beings, humans frequently infer other people’s thoughts and beliefs, a skill referred to as Theory of Mind (ToM). ToM enables effective social interactions, communication, and cooperation. Throughout development, the emergence of ToM has far-reaching consequences, being closely linked to children’s social-communicative skills^1,2^. Moreover, ToM capacity is related to prosocial behavior like sharing, positive relations with peers, cooperation, moral reasoning and emotions, and academic success^3^. A pivotal milestone in ToM development is robustly achieved by 4-5 years of age when children start to reason that others’ beliefs may differ from their own^4^. Despite this consistent developmental leap, there are important inter-individual variations in ToM performance in typically developing children^5,6^. In addition, delayed ToM development is characteristic of various cognitive disorders, including autism spectrum disorder (ASD), attention deficit hyperactivity disorder, developmental language disorders, and schizophrenia^7^. Despite this central role of ToM for human cognition and behavior, early predictors of individual ToM development are lacking. This has significantly hindered early identification and intervention. Being able to predict ToM, ideally before its emergence at preschool age, would have a crucial impact in cases of ToM deficiency.

A promising area in which to search for an early predictor of ToM development would be whole brain neural markers. Recent advances in computational modeling have enabled the identification of detailed networks relating to highly specific aspects of cognition, which can be subsequently used to predict behavior in other individuals^8–10^. This predictive modeling approach strands in contrast to correlational findings with individual brain structures, which do not typically allow prediction of individual differences in novel individuals. This is because, firstly, correlational studies tend to overfit the data, resulting in low generalizability to novel data^9^. To determine generalizable neural markers of ToM, it is therefore critical to use a cross-validated approach that tests identified brain-behavior relationships in an independent sample^9^. Secondly, there is growing consensus that brain regions do not work in isolation. Rather, interactions between multiple distributed brain areas and connections, creating networks, are the neural basis for cognitive functions^11^. Considering patterns of connectivity in the whole brain, referred to as the *connectome*, rather than individual brain regions or connections, has high potential for predicting interindividual differences in ToM development, even in individuals being tested for the first time^9^.

To attempt to identify generalizable neural markers of ToM development, we trained a cross-validated, connectome-based, predictive model (CPM^8^). CPMs are a powerful method to predict individual behavior from whole-brain patterns of connectivity^8,23^. They have been shown to predict various cognitive and affective functions, such as attention^10,12^, executive function^13,14^, and creativity^15,16^, and have successfully generated predictive models of atypical cognition^17–19^. Such predictive models are particularly relevant for development, where insights can be gained into how emerging brain networks give rise to cognition and behavior^19–22^.

Research on the neural basis of ToM in adults and older children (with established ToM abilities) has shown the involvement of a bilateral network comprising the temporoparietal junction (TPJ), the medial prefrontal cortex (mPFC), the precuneus (PC), the superior temporal sulcus (STS), and the inferior frontal gyri (IFG)^23,24–27^. However, this network matures relatively late in childhood^28–30^, and so far, no early neural predictors of ToM (i.e., before its onset in development) have been identified. Here, we reasoned that training a predictive neural model on a behavioral precursor of ToM may allow the identification of an early neural marker of subsequent ToM development. A promising candidate for a precursor of ToM in infancy is joint attention—a skill achieved towards the end of the first year of life when infants begin to coordinate their attention to items in their environment with other people^31^. Joint attention entails recognizing what other people attend to, and supports learning about other’s communicative and action goals^32–35^. As such, it critically shapes infants’ capacity to learn from others and their acquisition of social cognitive skills. Indeed, several longitudinal studies have shown positive behavioral correlations between joint attention and ToM^36–41^. The role of joint attention in early social cognitive development suggests that the neural maturation supporting joint attention in infancy may be predictive of later-developing ToM capacity. In adults, joint attention recruits brain areas partly overlapping with ToM regions (i.e., mPFC, pSTS, and TPJ)^42–51^, in addition to activations in the anterior insula, amygdala, and ventral striatum^48,51–55^. Functional Near Infrared Spectroscopy (NIRS) studies suggest activation of similar regions in infancy^20-23^ and correlations with connections between some of these regions were found^56,57^.

Identifying the neural footprint that predicts individual differences in joint attention development may thus serve as an early marker of ToM development. Here, we therefore set out to identify the brain networks supporting joint attention in infancy and hypothesized that these would be predictive of ToM abilities in preschool age. Leveraging the powerful modeling approach of CPM, with data from the Baby Connectome Project^58^, we first mapped joint attention scores to connectivity data. In brief, our model correlated 8- to 15-month-old infants’ behavioral joint attention scores with their resting-state fMRI whole-brain connectivity patterns and selected connections that significantly contributed to individual differences in behavior. These selected connections were summed to build a linear regression model that predicted behavioral joint attention scores from resting-state fMRI data of a different set of infants. Critically, in a next step, we showed that this model, trained on joint attention in infancy, was able to predict ToM performance in children aged 2 to 5 years.

## Results

### Connectome-based model significantly predicts joint attention scores

To investigate whether and which features of the resting-state connectome could predict infants’ joint attention scores, we randomly split the data into a training and a test set. A model was then built of the connections associated with joint attention in the training data set. We then tested this model in an independent subset of the data. The model generated on our training set (N = 97) significantly predicted joint attention scores from unseen rsfMRI connectivity in our independent test dataset (N = 47, Spearman rho = .329, p-value = .012, see Fig 1A).

**Figure 1.**
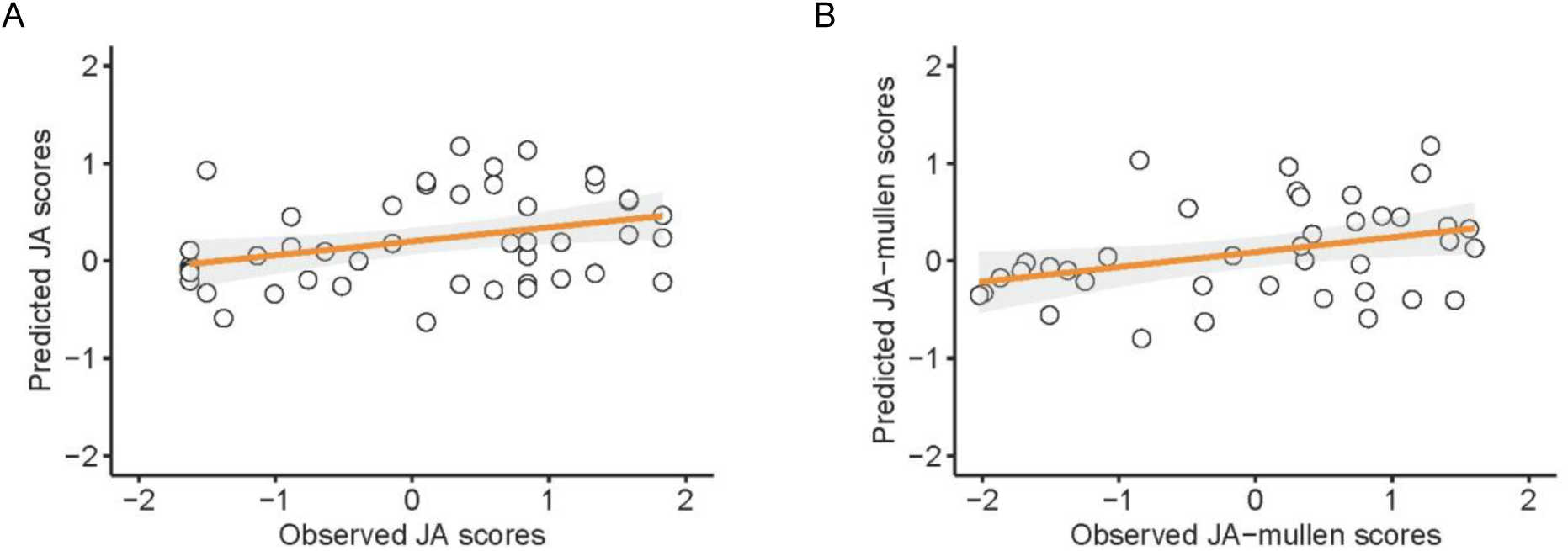
A connectome-based model built on joint attention data in a training dataset significantly predicted joint attention scores in an independent test data set of infants aged 8-15 months. A) Scatterplot of the correlation between predicted joint attention scores from the model and observed joint attention scores in the test data (N = 47, Spearman rho = .329, p = .012). B) Correlation of the joint attention model that controlled for general development by regressing out the *Mullen Scales of Early Learning* (B, N = 39, Spearman rho = .328, p = .029). All scores were z-normalized.

To test for the specificity of this prediction for joint attention beyond general development, we further controlled for the *Mullen Scales of Early Learning*. The resulting model was thus trained to predict joint attention, while controlling for general development. It also significantly predicted behavioral joint attention scores in the independent test data (N = 39, Spearman rho = .328, p-value = .029, see Fig 1B).

### Default mode and ventral-attention network connectivity predicts joint attention

To identify the network interactions that predicted joint attention, we computed network connectivity based on the 7-network parcellation^59^. In the predictive model trained on joint attention scores without controlling for general development, the majority of identified connections lay within sensory and motor networks (somatomotor (SMN) and visual (VIS)) and their interaction with the ventral attention network (VAN) and the default mode network (DMN) (details see Fig 2A/B). When controlling for infants’ general development, connections within the DMN and its interaction with the VAN became dominant, followed by connections between the DMN and SMN and within the VIS (details see Fig. 2C/D).

**Figure 2.**
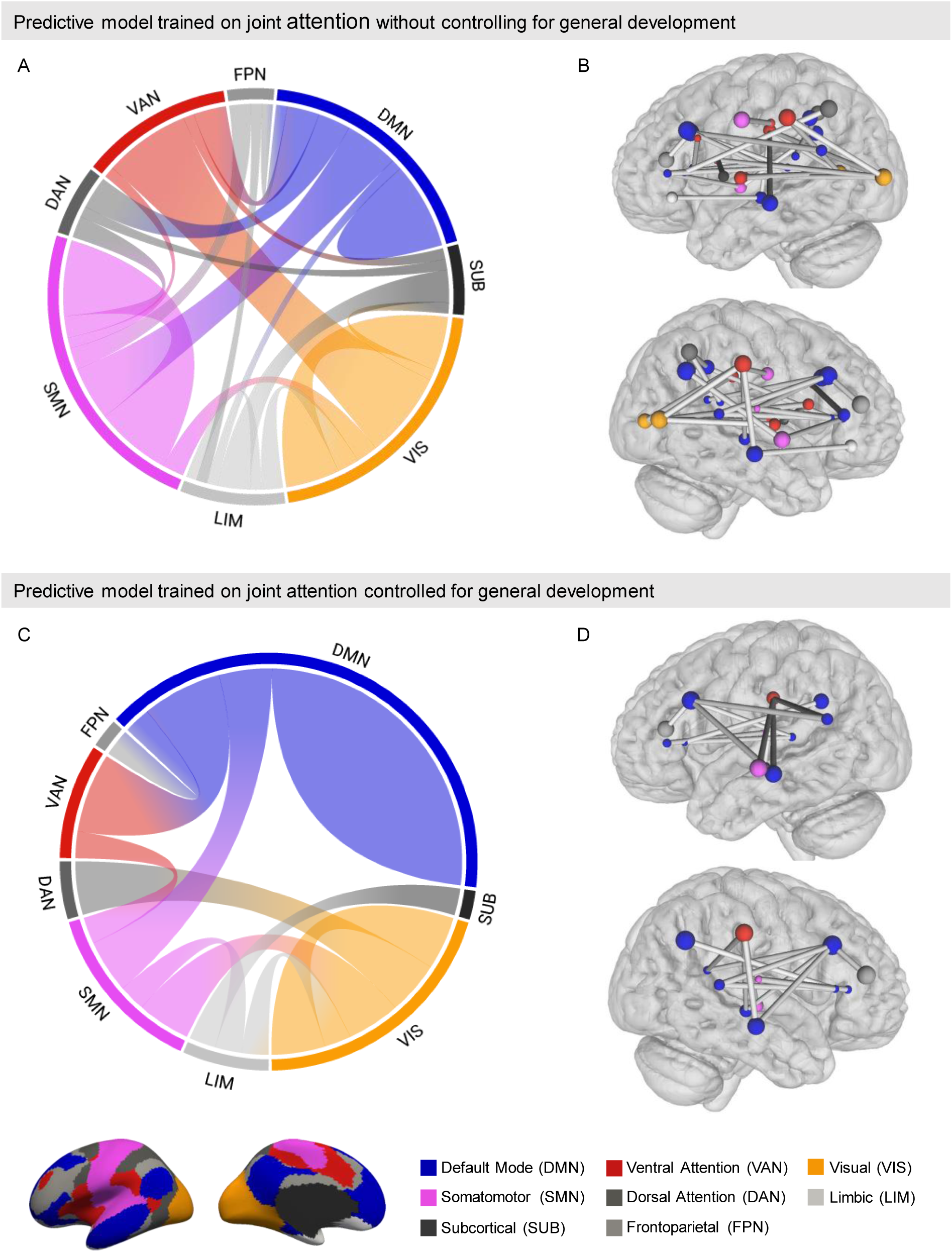
Network connections in the DMN and VAN predict joint attention. Shown are the connections between brain networks for A) the predictive model without controlling for general development and B) the model controlling for general development. Connections with nodes within the DMN and VAN for C) the model without controlling for general development and D) the model controlling for general development. Node size represents node degree, that is, the number of connections linked to this node. Network assignments are based on the Yeo 7-network parcellation (see color legend).

The connections that contributed to the joint attention model controlling for general development are listed, by network, in Table 1 (see Supplementary Table 1 for connections of the model without controlling for general development). In particular, within-DMN connections had nodes in the left banks of the superior temporal sulcus (bSTS), right rostral anterior cingulate cortex (rACC), right superior frontal gyrus (SFG), right inferior parietal gyrus (IPG), and bilateral middle temporal gyrus (MTG). Connections of the VAN with the DMN were between the right supramarginal gyrus (SMG) to the left bSTS and the bilateral MTG.

**Table 1.** List of connections between nodes contributing to the model that controlled for general development. Nodes are based on the UNC4D infant atlas, which uses the Desikan-Kiliany parcellation. Network assignments are based on the Yeo 7-network atlas. Abbreviations: DAN: dorsal attention network, DMN: default mode network, FPN: frontoparietal network, SMN: somatomotor network, SUB: subcortical, LIM: limbic network, VAN: ventral attention network, VIS: visual network; R: right and L: left hemispheres.

| Connection between |  |  | Node 2 |  |  |
| --- | --- | --- | --- | --- | --- |
| Node 1 |  |  |  |  |  |
| DMN | L | Bank superior temporal sulcus | DMN | R | Superior frontal gyrus |
| DMN | L | Middle temporal gyrus | DMN | R | Superior frontal gyrus |
| DMN | L | Bank superior temporal sulcus | DMN | R | Rostral anterior cingulate |
| DMN | L | Bank superior temporal sulcus | DMN | R | Superior frontal gyrus |
| DMN | R | Inferior parietal gyrus | DMN | R | Rostral anterior cingulate |
| DMN | R | Middle temporal gyrus | DMN | R | Superior frontal gyrus |
| DMN | L | Bank superior temporal sulcus | VAN | R | Supra marginal gyrus |
| DMN | L | Middle temporal gyrus | VAN | R | Supra marginal gyrus |
| DMN | R | Middle temporal gyrus | VAN | R | Supra marginal gyrus |
| DMN | L | Rostral anterior cingulate | SMN | R | TransverseTemp |
| DMN | R | Rostral anterior cingulate | SMN | R | TransverseTemp |
| VIS | R | Pericalcarine cortex | DAN | L | Superior parietal gyrus |
| VIS | R | Pericalcarine cortex | DAN | R | Superior parietal gyrus |
| VIS | R | Fusiform gyrus | VIS | R | Lingual gyrus |
| VIS | R | Fusiform gyrus | VIS | R | Pericalcarine cortex |
| VIS | L | Lateral occipital cortex | SMN | R | Superior temporal gyrus |
| VIS | L | Lateral occipital cortex | SMN | R | TransverseTemp |
| VAN | R | Supra marginal gyrus | SMN | L | Superior temporal gyrus |
| LIM | L | Entorhinal cortex | SMN | R | Paracentral gyrus |
| LIM | L | Frontal pole | SUB | R | Caudate |
| LIM | R | Entorhinal cortex | VIS | R | Fusiform gyrus |
| DMN | R | Superior frontal gyrus | FPN | R | Rostral middle front gyrus |

### Joint attention model in infancy predicts later Theory of Mind at preschool age

Critically, we then tested whether this connectome identified for joint attention in infancy would predict later-developing ToM in the preschool years. To this aim, we applied the joint attention model that controlled for general development to the resting-state data of preschool children aged 2 to 5 years (N = 37). Notably, the joint attention model, which was initially trained in infants aged 8 to 15 months, significantly predicted the emergence of ToM reasoning in the independent sample of preschool-aged children aged 2 to 5 years (N = 37, rho = .336, *p* = .021, Fig 3).

**Figure 3.**
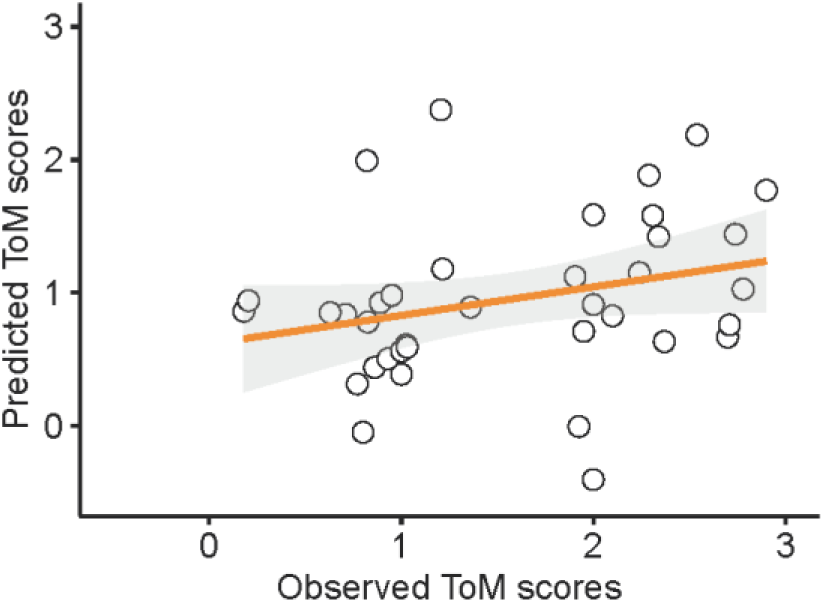
A connectome-based model built on joint attention data in infancy significantly predicted ToM capacity in preschool children. Scatterplot of the correlation between predicted ToM scores from the model and observed ToM scores (N = 37, rho = .336, p = .021).

## Discussion

The emergence of ToM in cognitive development marks a pivotal milestone, robustly achieved in preschool age. Despite its importance for human cognition, there are no known early predictors of individual ToM development. We reasoned that the neural footprint identified from a potential behavioral precursor of ToM in infancy—joint attention—may provide such an early marker. To this end, we modeled 8- to 15-month-old infants’ whole brain connectivity with connectome-based predictive modeling to identify network patterns that predicted engagement in joint attention, while controlling for general cognitive development. Indeed, this model significantly predicted joint attention scores, also in an independent cohort of individuals. In other words, our model was able to predict the behavioral performance of a new group of infants merely from the patterns of their resting state fMRI data. Critically, this same model successfully predicted the development of ToM in children several years older. That is, the model, which identified a pattern of connectivity predictive of joint attention in multiple cohorts of infants, was able to predict ToM abilities of a third cohort of much older children.

The predictive power of our model likely rests on the cross-validated, whole-connectome, predictive modelling approach that allowed the discovery of whole-brain network patterns predictive of joint attention, rather than being limited to individual connections or brain regions. As the networks identified for joint attention also predicted ToM reasoning in older children, these findings suggest a causal relation in which the maturation of these networks drives ToM development. This is in line with the proposed role of joint attention as a catalyst for social cognitive development and ToM^31^ and with partly overlapping brain regions observed for joint attention and ToM^60^. The dominant connections in the model lay within the DMN, linked to various domains of higher-order cognition and largely overlapping with the network activated for ToM^61,62^. This supports the notion that the DMN’s capacity to integrate sensory signals and external attention^63^ within social contexts is already in place in infancy. The DMN was previously shown to be active during real-time synchronization in social interactions^64^, allowing a mutual attunement to one another, which is highly relevant for infants and their caregivers^65^. In particular, during moments of mutual gaze, smiling, and joint attention, local neural couplings in infants and caregivers were observed in the mPFC, a core region of the DMN^66^. Moreover, the emergence of mature ToM in the preschool years was associated with increased structural connectivity of the DMN and its connection to regions involved in language and executive function^29^.

Next to within-DMN connectivity, its interaction with the VAN, followed by sensory network interactions, contributed to the predictive model. This interplay of sensory input processed in the VIS and SMN and information filtering in the VAN may contribute to the integration of sensory input (e.g., the others’ gaze or pointing and target of attention), attentional control (to focus on the attentionally guiding cues), and coordination of one’s movements (to follow these cues) required for joint attention. Thereby, bottom-up, stimulus-driven processing in the VAN may allow infants to attune and react to striking and attention-grabbing cues in their environment. For example, the gaze cues of another person and the highlighted objects are particularly salient, and therefore facilitate infants learning in social interactions in a bottom-up fashion^67–69^. An initial model of joint attention relied on connections within sensory networks and their interactions with the DMN and VAN. However, when accounting for children’s general cognitive development, within-DMN connectivity became the dominant predictive contributor, followed by its interaction with the VAN. Connections within the somatomotor network, in turn, diminished. This indicates that these connections likely pertain to general developmental processes, such as motor coordination, that may facilitate joint attention but lack specificity to it. This interplay of higher-order, attention, and sensory networks may contribute to infants’ development beyond joint attention, giving rise to the 9-month revolution^31^, a period of developmental leaps across different social cognitive domains. Indeed, our results suggest that these networks already play a foundational role in the emergence of joint attention, which then, in turn, sets the stage for later-developing ToM.

While joint attention is a directly perceivable activity, the later-developing skill of ToM is used to infer *invisible* mental states, that is, content which is decoupled from perceptual input. Our finding that the networks predictive of joint attention also predicted later ToM reasoning fit with the recent suggestions that the interaction between bottom-up networks and the DMN may contribute to decoupling content from perceptual input^70,71^. This possible shift from perceptual access to higher, perceptually decoupled cognition is reflected in the maturational trajectories of brain networks. Indeed, sensory networks mature first, with adult-like functional patterns by 9 months, while attention networks and the DMN mature later^72,73^. In line with this idea, a study correlating behavioral joint attention scores with functional connectivity found that 24-month-old infants showed decreased coupling between sensory networks and higher-order networks compared to 12-month-olds^56^.

Given its predictive power for ToM and joint attention, our model and the employed CPM approach has high potential to provide early neuro-markers of atypical development with impairments in ToM and joint attention, such as in autism spectrum disorder (ASD)^74,75^. Because of its function as a catalyst, joint attention impairments at this early stage in life have far-reaching and negatively cascading consequences^38,41^. Thus, early diagnosis and intervention are of great importance. Indeed, imaging-based models have been used to predict ASD^76,77^ and ADHD^78^. Our connectome-based approach, specifically modeling joint attention and predicting ToM, could extend the diagnostic power of such models. Applying our model to these disorders could lead to early interventions targeting the contributing brain networks predictive of joint attention and ToM.

In conclusion, we have demonstrated that a predictive model of joint attention, trained on the whole-brain connectome of 8- to 15-month-old infants, significantly predicted ToM capacity in children several years later. Dominant connections of the model lay within the DMN and its interaction with the VAN. We propose that the development of joint attention is underpinned by increased connectivity of the DMN and its coupling to bottom-up processing in the VAN, bridging the transition from perceptually-based processing to more abstract social cognitive processes that are decoupled from perception, such as ToM. The possibility to predict ToM at this early stage in development reveals intriguing potential for the early diagnosis of social cognitive disorders like ASD, ADHD, and delayed language development. An exciting avenue for future research is whether a similar approach can predict other cognitive developments linked to joint attention, such as language and other social cognitive functions.

## Methods

### Baby Connectome Project Dataset

For this study, we used open-access data from the Baby Connectome Project (BCP^58^), available via the NIMH Data Archive (ID: 2848). We were interested in participants with resting state data and behavioral assessments of either joint attention or theory of mind. Our joint attention analysis included N = 92 infants (51 female) aged 8 to 15 months (median 11 months) with a total of 144 scans. Furthermore, for the prediction of Theory of Mind scores in a different cohort of preschool-aged children, N = 37 children (21 female) aged 24 to 60 months (median 34 months) with a total of 38 scans were included.

### Behavioral Assessment

#### Joint Attention

Joint attention was assessed with the *Dimensional Joint Attention Assessment*^57^ in infants aged 8 to 15 months. Several novel toys were distributed in a room. The experimenter sat in front of the infant and initiated four joint attention prompts with decreasing difficulty. First, the experimenter shifted their gaze and head toward a toy. Secondly, the experimenter additionally said, “Look at that”. Then, the experimenter shifted their gaze and head and pointed at the toy. Last, the experimenter used all cues of shifting gaze and head, pointing, and verbalization. If the infant responded to one prompt, the series was accomplished, and the infant was allowed to play with the toy. If the infant did not respond, the experimenter continued with the next level of difficulty. These maximal four prompts were repeated with four different toys. For each series, the infant was scored from 0 (no response) to 4 (responding to the first, most difficult prompt), and the mean of all series was used for further analysis. Joint attention assessment and MRI data acquisition occurred within a timespan of two weeks.

#### General Development

The general development of infants was assessed with the *Mullen Scales of Early Learning*^79^. The scale spans the domains of gross motor, fine motor, visual reception, receptive language, and expressive language. The scale comprises interactive tasks and was administered in a standardized format. For our analysis, an aggregated score across all subdomains was used.

#### Theory of Mind

Theory of Mind skills in a different cohort of preschool-aged children was assessed with the *Children’s Social Understanding Scale* (CSUS^80^, which is a parent-report questionnaire that was cross-validated in several studies with children’s performance in different ToM tasks. The scale comprises evaluating various aspects of Theory of Mind, including children’s understanding of other people’s beliefs, knowledge, perceptual access, desires, intentions, and emotions. Theory of Mind and MRI data were acquired within one month.

### MRI Data Acquisition

Infants under 3 years of age were scanned during natural sleep (for a review see^81^, for detailed procedures^58^). Parents of children above 3 years could choose between sleep and awake MRI. All images were acquired on a Siemens 3T Prisma with a 32-channel head coil. Sequences used in the BCP were specifically tailored to the needs of infant MRI (e.g., by using quieter and shorter sequences). Anatomical MRI was acquired with MPRAGE (0.8 mm isotropic, TE=2.24 ms, TR=2400/1060 ms, 6:38 minutes). Resting-state fMRI was acquired with a single-shot EPI sequence (2 mm isotropic, TE=37 ms, TR=800 ms, 5:47 min). For further details, see the BCP acquisition pipeline^58^.

### MRI Data Preprocessing

MRI data from the BCP were preprocessed using the state-of-the-art infant-specific pipeline niBabies 22.0.1, derived from fMRIPrep^82^ with the UNC 4D volumetric atlas^83^ in *MNIInfant* space. Preprocessing was performed with the respective age-group-specific template to account for age-related brain size differences. Preprocessed data were quality controlled by checking for proper brain extraction, T1w/T2w registration and quality of functional data. For details, see supplements. For additional details, see Supplementary Methods 1 for pipeline-generated boilerplate.

### Connectome Construction

To obtain connectivity matrices from the preprocessed rsfMRI data, CONN toolbox 21.a (RRID:SCR_009550^84^) was used. As suggested, an additional functional smoothing step with a Gaussian kernel of 6 mm full-width half maximum (FWHM)^85^ was performed after importing the preprocessed rsfMRI data. Furthermore, CONN’s default denoising pipeline, including linear regression of potential confounding effects in the BOLD signal and temporal bandpass filtering, was performed. ROI-to-ROI connectivity was calculated with the UNC-4D^83^ infant atlas containing a cortical parcellation of 82 regions based on the Desikan-Killiany parcellation^86^. For additional details, see Supplementary Methods 2 for pipeline-generated boilerplate.

### Connectome Predictive Modeling

#### Building a model predictive of joint attention in the training data

Connectome predictive modeling (CPM) was used to predict behavioral JA scores from resting-state connectivity matrices. CPM involves the steps of feature selection, model building, and model validation described below (for details see^8^). We randomly split our data into a training (N = 97) and test (N = 47) set to obtain independent datasets. In the first step of CPM, connections whose individual variations could explain differences in behavior were obtained. To achieve this, we correlated joint attention scores with the edges (functional connections) of rfMRI connectivity matrices in the training set. Edges above a correlation threshold of p = 0.005 were selected for the model. We used a leave-one-out cross-validation approach, repeating the edge selection step 97 times while always leaving out one participant. The final set of edges that make up the joint attention network was based on edges found in 90% of all leave-one-out iterations. Each participant’s connectivity strength was summed for the selected edges, and this summed network strength was used to build linear regressions that model the relation between connectivity strength and joint attention.

#### Predicting joint attention scores and validating the model

The built joint attention model was validated by applying the model to the unseen, independent test dataset (N = 47). In the test dataset, individual network strength for the selected joint attention edges was computed as described above. Then, joint attention scores were predicted based on resting-state connectivity, and the predictive performance of the model was evaluated with a one-sided Spearman correlation between the predicted scores and the measured joint attention scores.

#### Controlling for general development

To ensure the specificity of our joint attention model, we controlled for the general development of children. For this, we built a model as described above in the training data but regressed out the *Mullen Scales of Early Learning* from the joint attention scores (N = 79). The model was validated in the independent test data as described above for the uncontrolled model (N = 39).

#### Predicting Theory of Mind scores

To test whether the built models predict ToM scores in older children (N = 37), we applied the model predictive of joint attention and controlled for general development on the resting-state data of children with ToM measures to predict ToM scores. Model performance was assessed with a Spearman correlation between the predicted and the actual ToM scores.

#### Optimizing the Feature Selection Threshold

As suggested by Shen et al.^8^, we determined the optimal feature selection threshold that determines which edges are considered in the model. For this, we split the original dataset into a training (N=97) and test (N=47) set. In our training dataset, we tested four different, commonly used, feature selection thresholds (p = 0.05, p = 0.01, p = 0.005, p = 0.001) with a leave-one-out cross-validation approach. We evaluated each model built with a different feature selection threshold by comparing their predictive performance (Spearman rho). For all analyses in the test set, we used the feature selection threshold of p = 0.005 obtained from the training dataset. See performance of the tested feature selection thresholds in Supplementary Table 2.

#### Controlling for Motion

As suggested by Shen et al.^8^, we controlled for motion by correlating motion parameters (framewise displacement according to Power) with behavior and resting state connectivity. Neither a correlation of motion with behavior (Spearman rho=.017, p=.825) nor a prediction of motion by the predictive model (Spearman rho=-.133, p=.372) was found.

#### Network Connectivity

Brain network connectivity was investigated by using the mapping of the 7-network atlas^59^ and the Desikan-Kiliany parcellation, on which the UNC 4D infant atlas^83^ is based. This mapping between different atlases was computed with spatial permutations^87^. We assigned each node in the UNC 4D atlas to one of the seven networks based on its highest overlap. For network connectivity, the sum of connections between networks was computed for visualization.

## Acknowledgements

Data used in this study were provided by the Baby Connectome Project, and research initiative of the Neuroscience Blueprint. This research was funded by a scholarship of the FAZIT Stiftung to CS, and grants by the German Research Foundation (DFG) to PB (project number FR 519/20-2) and to CGW (project number GR 5421/1-2).

## Author contributions

Conceptualization – CS, CGW, and PB. Data curation, Formal analysis, Methodology, Validation, and Visualization – CS. Supervision – CGW. Writing, original draft – CS. Writing, review and editing – CGW and PB.

