## Supplements for "Predicting Theory of Mind in children from the infant connectome"

**Supplementary Table 1 List of connections between nodes contributing to the model without controlling for general development**. Nodes are based on the UNC4D infant atlas, which uses the Desikan-Kiliany parcellation. Network assignments are based on the Yeo 7-network atlas. Abbreviations: DAN: dorsal attention network, DMN: default mode network, FPN: frontoparietal network, SMN: somatomotor network, SUB: subcortical, LIM: limbic network, VAN: ventral attention network, VIS: visual network; R: right and L: left hemispheres.

| **Connection between** | | | | | |
|---|---|---|---|---|---|
| **Node 1** | | | **Node 2** | | |
| VIS | L | Lateral occipital cortex | VAN | L | Supra marginal gyrus |
| VIS | L | Lateral occipital cortex | VAN | R | Pars opercularis |
| VIS | L | Lateral occipital cortex | VAN | R | Supra marginal gyrus |
| VIS | L | Lateral occipital cortex | VAN | R | Insula |
| VIS | R | Lateral occipital cortex | VAN | R | Supra marginal gyrus |
| VIS | R | Lateral occipital cortex | VAN | R | Insula |
| VIS | R | Lateral occipital cortex | VAN | L | Supra marginal gyrus |
| VIS | R | Lateral occipital cortex | VAN | L | IsthmusInsula |
| SMN | L | Paracentral gyrus | SMN | R | Paracentral gyrus |
| SMN | L | Paracentral gyrus | SMN | R | Postcentral gyrus |
| SMN | L | Paracentral gyrus | SMN | R | Precentral gyrus |
| SMN | L | Postcentral gyrus | SMN | R | Postcentral gyrus |
| SMN | L | Postcentral gyrus | SMN | R | Precentral gyrus |
| SMN | L | Precentral gyrus | SMN | R | Paracentral gyrus |
| SMN | L | Precentral gyrus | SMN | R | Precentral gyrus |
| VIS | L | Fusiform gyrus | VIS | R | Cuneus |
| VIS | L | Fusiform gyrus | VIS | R | Lingual gyrus |
| VIS | L | Fusiform gyrus | VIS | R | Pericalcarine cortex |
| VIS | L | Lateral occipital cortex | VIS | R | Pericalcarine cortex |
| VIS | R | Fusiform gyrus | VIS | R | Lingual gyrus |
| DMN | L | Rostral anterior cingulate | SMN | R | Superior temporal gyrus |
| DMN | L | Rostral anterior cingulate | SMN | R | TransverseTemp |
| DMN | L | Superior frontal gyrus | SMN | R | TransverseTemp |
| DMN | R | Inferior parietal gyrus | SMN | R | Superior temporal gyrus |
| DMN | R | Rostral anterior cingulate | SMN | R | TransverseTemp |
| DMN | L | Bank superior temporal sulcus | DMN | R | Superior frontal gyrus |
| DMN | L | Rostral anterior cingulate | DMN | L | Bank superior temporal sulcus |
| DMN | L | Bank superior temporal sulcus | DMN | R | Rostral anterior cingulate |
| DMN | L | Bank superior temporal sulcus | DMN | R | Superior frontal gyrus |
| DMN | L | Isthmus cingulate | DAN | R | Superior parietal gyrus |
| DMN | L | Precuneus | DAN | R | Superior parietal gyrus |
| DMN | R | Rostral anterior cingulate | DAN | L | Superior parietal gyrus |
| DMN | L | Middle temporal gyrus | VAN | R | Supra marginal gyrus |
| DMN | R | Middle temporal gyrus | VAN | R | Supra marginal gyrus |
| DMN | R | ParsTriangularis | VAN | L | Caudal anterior cingulate |
| LIM | L | Frontal pole | SUB | L | Caudate |
| LIM | L | Frontal pole | SUB | R | Caudate |
| LIM | L | Frontal pole | SUB | R | Pallidum |
| VIS | L | Lateral occipital cortex | SMN | R | TransverseTemp |
| VIS | L | Lingual gyrus | SMN | L | Superior temporal gyrus |
| SMN | L | Postcentral gyrus | DAN | R | Superior parietal gyrus |
| SMN | R | Postcentral gyrus | DAN | L | Superior parietal gyrus |
| FPN | R | Caudal middle frontal gyrus | SMN | L | Superior temporal gyrus |
| FPN | R | Caudal middle frontal gyrus | SMN | R | Superior temporal gyrus |

| | | | | | | | |
|---|---|---|---|---|---|---|---|
| | FPN | R | Corpus Callosum | | LIM | R | Entorhinal cortex |
| | FPN | R | Rostral middle front gyrus | | DMN | R | Superior frontal gyrus |
| | LIM | L | Entorhinal cortex | | VIS | R | Fusiform gyrus |
| | LIM | L | Entorhinal cortex | | LIM | R | Medial orbitofrontal gyrus |
| | LIM | R | Entorhinal cortex | | VIS | R | Fusiform gyrus |
| | LIM | R | Lateral orbitofrontal gyrus | | DMN | R | Middle temporal gyrus |
| | SMN | L | Precentral gyrus | | VAN | R | Posterior cingulate |
| | SUB | L | Thalamus | | DAN | R | Superior parietal gyrus |
| | SUB | L | Putamen | | VAN | R | Pars opercularis |
| | VIS | R | Pericalcarine cortex | | SUB | R | Hippocampus |

**Supplementary Table 2 Optimization of the feature selection threshold.** We determined the optimal feature selection threshold that determines which edges are considered in the model. In our training dataset, we tested four different, commonly used, feature selection thresholds (p=0.05, p=0.01, p=0.005, p=0.001) with a leave-one-out cross-validation approach. For all analyses in the test set, we've used the feature selection threshold of p=0.005 obtained from the training data.

| Feature selection threshold | Spearman's rho | p-value |
|---|---|---|
| 0.05 | 0.2946 | 0.0034 |
| 0.01 | 0.3279 | 0.0012 |
| 0.005 | 0.4103 | 0.00003 |
| 0.001 | 0.3243 | 0.0012 |

**Supplementary Methods 1 Boilerplate from preprocessing pipeline niBabies 22.0.1**

*Preprocessing of B0 inhomogeneity mappings*

A B0-nonuniformity map (or field map) was estimated based on two echo-planar imaging (EPI) references with FSL topup.

*Anatomical data preprocessing*

All of the T1-weighted images were corrected for intensity nonuniformity (INU) with `N4BiasFieldCorrection`, distributed with ANTs 2.3.3. The T1w reference was then skull-stripped with a modified implementation of the `antsBrainExtraction.sh` workflow (from ANTs), using the UNC4Datlas as the target template. An anatomical reference map was computed after registration of INU-correction images using FreeSurfer `mri_robust_template`. Brain tissue segmentation of cerebrospinal fluid (CSF), white matter (WM) and gray matter (GM) was performed on the brain-extracted T1w using FSL FAST. Volume-based spatial normalization to one standard space (age-specific UNC4D atlas) was performed through nonlinear registration with `antsRegistration` (ANTs 2.3.3) using brain-extracted versions of both the T1w reference and the T1w template.

*Functional data preprocessing*

For each of the BOLD runs, the following preprocessing was performed. Head-motion parameters with respect to the BOLD reference (transformation matrices and six corresponding rotation and translation parameters) are estimated before any spatiotemporal filtering using FSL `mcflirt`. The estimated field map was then aligned with rigid registration to the target EPI (echo-planar imaging) reference run. The field coefficients were mapped onto the reference EPI using the transform. BOLD runs were slice-time corrected to the midpoint of 0.5 of slice acquisition range using AFNI `3dTshift` from AFNI. The BOLD reference was then coregistered to the T1w reference using FSL `flirt` with the boundary-based registration cost function. Coregistration was configured with nine degrees of freedom to account for distortions remaining in the BOLD reference. First, a reference volume and its skull-stripped version were generated using a custom methodology of fMRIPrep. Several confounding time series were calculated based on the preprocessed BOLD: framewise displacement (FD), DVARS and three regionwise global signals. FD was computed using two formulations following Power (absolute sum of relative motions) and Jenkinson (relative root mean square displacement between affines). FD and DVARS are calculated for each functional run, both using their implementations in Nipype. The three global signals are extracted within the CSF, the WM, and the whole-brain masks. Additionally, a set of physiological regressors were extracted to allow for component-based noise correction. Principal components are estimated after high-pass filtering of the preprocessed BOLD time series (using a discrete cosine filter with a 128 s cutoff) for the two CompCor variants: temporal (tCompCor) and anatomical (aCompCor). tCompCor components are then calculated from the top 2% variable voxels within the brain mask. For aCompCor, three probabilistic masks (CSF, WM and combined CSF+WM) are generated in anatomical space. The implementation differs from that of Behzadi et al. in that instead of eroding the masks by 2 pixels on BOLD space, the aCompCor masks are subtracted from a mask of pixels that likely contain a volume fraction of GM. This mask is obtained by thresholding the corresponding partial volume map at 0.05, and it ensures that components are not extracted from voxels containing a minimal fraction of GM. Finally, these masks are resampled into BOLD space and binarized by thresholding at 0.99 (as in the original implementation). Components are also calculated separately within the WM and CSF masks. For each CompCor decomposition, the k components with the largest singular values are retained, such that the retained components' time series are sufficient to explain 50 percent of the variance across the nuisance mask (CSF, WM, combined, or temporal). The remaining components are dropped from consideration. The head-motion estimates calculated in the correction step were also placed within the corresponding confounds file. The confounding time series derived from head motion estimates and global signals were expanded with the inclusion of temporal derivatives and quadratic terms for each. Frames that exceeded a threshold of 0.5 mm FD or 1.5 standardized DVARS were annotated as motion outliers. The BOLD time series were resampled into standard space, generating a preprocessed BOLD run in UNC4D space. First, a reference volume and its skull-stripped version were generated using a custom methodology of fMRIPrep. All resamplings can be performed with a single interpolation step by composing all the pertinent transformations (i.e., head-motion transform matrices, susceptibility distortion correction when available, and coregistrations to anatomical and output spaces). Gridded (volumetric) resamplings were performed using `antsApplyTransforms` (ANTs), configured with Lanczos interpolation to minimize the smoothing effects of other kernels. Nongridded (surface) resamplings were performed using `mri_vol2surf`(FreeSurfer).

**Supplementary Methods 2 Boilerplate from preprocessing pipeline CONN**

Results included in this manuscript come from analyses performed using CONN (RRID:SCR_009550) release 21.a and SPM (RRID:SCR_007037) release 12.7771.

*Denoising*

In addition, functional data were denoised using a standard denoising pipeline including the regression of potential confounding effects characterized by white matter timeseries (5 CompCor noise components), CSF timeseries (5 CompCor noise components), motion parameters and their first order derivatives (12 factors), outlier scans (below 334 factors), session and task effects and their first order derivatives (2 factors), and linear trends (2 factors) within each functional run, followed by bandpass frequency filtering of the BOLD timeseries between 0.008 Hz and 0.09 Hz. CompCor noise components within white matter and CSF were estimated by computing the average BOLD signal as well as the largest principal components orthogonal to the BOLD average, motion parameters, and outlier scans within each subject's eroded segmentation masks.

*ROI-to-ROI connectivity*

ROI-to-ROI connectivity matrices were estimated characterizing the functional connectivity between each pair of regions among 82 ROIs. Functional connectivity strength was represented by Fisher-transformed bivariate correlation coefficients from a general linear model (weighted-GLM), estimated separately for each pair of ROIs, characterizing the association between their BOLD signal timeseries. In order to compensate for possible transient magnetization effects at the beginning of each run, individual scans were weighted by a step function convolved with an SPM canonical hemodynamic response function and rectified
